# DNA unwinding mechanism of a eukaryotic replicative CMG helicase

**DOI:** 10.1101/737015

**Authors:** Zuanning Yuan, Roxana Georgescu, Lin Bai, Dan Zhang, Huilin Li, Mike O’Donnell

## Abstract

High-resolution structures have not been reported for replicative helicases at a replication fork at atomic resolution, a prerequisite to understand the unwinding mechanism. The eukaryotic replicative CMG helicase contains a Mcm2-7 motor ring, with the N-tier ring in front and the C-tier motor ring behind. The N-tier ring is structurally divided into a zinc finger (ZF) sub-ring followed by the OB fold ring. Here we report the cryo-EM structure of CMG on forked DNA at 3.9 Å, revealing that parental DNA enters the ZF sub-ring and strand separation occurs at the bottom of the ZF sub-ring, where the lagging strand is blocked and diverted sideways by OB hairpin-loops of Mcm3, Mcm4, Mcm6, and Mcm7. Thus, instead of employing a separation pin, unwinding is achieved via a “dam-and-diversion tunnel” for steric exclusion unwinding. The C-tier motor ring contains spirally configured PS1 and H2I loops of Mcms 2, 3, 5, 6 that translocate on the spirally-configured leading strand, and thereby pull the preceding DNA segment through the diversion tunnel for strand separation.

## INTRODUCTION

Replicative helicases of all cell types are hexameric rings composed of a N-tier ring of the NTD domains and a C-tier ring (O’Donnell and Li, 2018). The 11-subunit eukaryotic helicase CMG (Cdc45, Mcm2-7, GINS) contains a hexameric Mcm2-7 motor ring (Enemark and Joshua-Tor, 2008; Ilves et al., 2010; O’Donnell and Li, 2018; Parker et al., 2017). The NTD ring of the Mcm2-7 motor is further divided into an upper sub-ring of six zinc finger (ZF) domains and a lower sub-ring composed of six oligosaccharide/oligonucleotide-binding (OB) domains that bind single stranded DNA (ssDNA) (Li and O’Donnell, 2018). The C-tier contains the motors with ATP sites located at subunit interfaces (Davey et al., 2003; Lyubimov et al., 2011). The ATP binding site of bacterial helicases is based on the RecA fold, while eukaryotic helicase ATP sites are based on the AAA+ fold (Erzberger and Berger, 2006; Li and O’Donnell, 2018; Lyubimov et al., 2011; O’Donnell and Li, 2018). While bacterial replicative helicases travel 5’-3’ with the C-tier motors leading the way, eukaryotic helicases track 3’-5’ on DNA with the N-face in front, pushed by motors in the C-tier (Enemark and Joshua-Tor, 2008; Li and O’Donnell, 2018; O’Donnell and Li, 2018). The way in which helicases engage ssDNA in the motor domains has been documented for several different hexameric replicative helicases and involve binding to loops in the ATPase domains (Abid Ali et al., 2017; Enemark and Joshua-Tor, 2006; Gao et al., 2019; Itsathitphaisarn et al., 2012; Li and O’Donnell, 2018; Meagher et al., 2019; O’Donnell and Li, 2018; Skordalakes and Berger, 2006; Thomsen and Berger, 2009). In eukaryotes the main loop is referred to as PS1 (e.g. BPV viral E1 and SV40 T-antigen of superfamily 4 helicases); while archaeal MCMs and eukaryotic Mcm2-7 of CMG (superfamily 6 helicases) contain the pre-sensor 1 (PS1) loop and also a helix 2 insertion (H2I) loop of unknown function. The PS1 loops are proposed to pull the ssDNA such that the parental duplex splits at the top of the helicase, a process known as the steric exclusion mechanism of helicase action, because one strand is excluded from the central channel while the other strand is pulled through the central channel (Fu et al., 2011). In simple terms, the helicases act as a moving wedge to separate the strands of dsDNA. However, CMG has a wide central channel and easily accepts dsDNA (Langston et al., 2017; Wasserman et al., 2019). Hence details of how steric exclusion actually splits the strands rather than bringing them both into the central channel of the helicase are unknown, as high-resolution structures of helicases with bound forked DNA are lacking.

In a recently reported cryo-EM structure of the T7 replisome complexed with a forked DNA, only ssDNA enters the helicase ring at the C-tier, with no interaction between the helicase and the parental duplex (Gao et al., 2019). Interestingly, the dsDNA and ssDNA orient perpendicularly to one another, and this geometry is hypothesized to greatly accelerate the rate of T7 helicase activity (Gao et al., 2019). The CMG replicative helicase of eukaryotic cells contains 11 subunits in which the Mcm2-7 forms an ATPase ring for DNA unwinding (Costa et al., 2014; Ilves et al., 2010; Moyer et al., 2006; Sun et al, 2015; Yuan et al., 2016). Our earlier study of the budding yeast Saccharomyces cerevisiae CMG bound to a forked DNA demonstrated that the parental duplex enters the N-tier of CMG a short distance (Georgescu et al., 2017), which was different from the classic steric exclusion models (Enemark and Joshua-Tor, 2008; Lyubimov et al., 2011). Unlike the T7 replisome structure (Gao et al., 2019), the eukaryotic CMG-forked DNA structure showed a nearly in-line orientation of dsDNA and unwound leading ssDNA, having only about a 17° angle between them (Georgescu et al., 2017). Moreover, another cryo-EM structure of CMG-Pol ε−forked DNA showed that dsDNA enters inline with the central channel (Goswami et al., 2018). However, Pol ε binds on the C-tier bottom surface (Sun et al., 2015), and might not be expected to alter DNA geometry entering the N-tier. Thus It is currently unclear if protein factors interacting with the N-tier surface of the Mcm2-7 ring, such as Mcm10, would alter the entry angle of the parental dsDNA into a perpendicular configuration for more rapid unwinding, as observed when Mcm10 is bound to CMG (Langston et al., 2017; Looke et al., 2017). One motivation of the current work is to examine this question.

In the T7 replisome, the split point of the DNA duplex is outside the helicase and therefore the forked DNA region is not well stabilized for high resolution visualization (Gao et al., 2019). Nevertheless, a β-hairpin in the DNA polymerase, outside the helicase, is proposed to function as the separation pin for dsDNA unwinding. In contrast, the fork junction is inside the chamber in the eukaryotic CMG helicase. We previously used a dual streptavidin block on DNA to stall the CMG unwinding at a fork in the presence of ATP and determined a medium-resolution cryo-EM structure of CMG on forked DNA (Georgescu et al., 2017). While this study revealed the N-first orientation of CMG during DNA tracking, the 6.7-Å resolution was too low to resolve the key structural features critical for DNA unwinding (Georgescu et al., 2017). In the cryo-EM study of the DNA fork-containing CMG in the presence of ATPγS and Pol ε, the average resolution reached to about 5 Å (Goswami et al., 2018). However, there was a density gap between the parental dsDNA region in the N-tier and the leading strand DNA in the C-tier motor ring; in other words, the DNA fork junction region was also not visualized in that study. Therefore, the primary motivation of this manuscript is to visualize the forked DNA junction structure at atomic or near atomic resolution−high enough resolution to define a precise mechanism of dsDNA unwinding by a eukaryotic helicase performing steric exclusion, that could otherwise draw both DNA strands through the central channel.

## RESULTS

### The 3.9 Å structure of CMG − forked DNA in the presence of Mcm10

Mcm10 binds tightly to CMG (Langston et al., 2017; Mayle et al., 2019), enhances CMG binding to forked DNA (Wasserman et al., 2019), and stimulates the rate and processivity of CMG (Langston et al., 2017). Therefore, in the current work, we assembled CMG on the double-streptavidin stalled forked DNA in the presence of Mcm10 and carried out cryo-EM analysis of the assembled product. We found that Mcm10 increased the percentage of CMG-forked DNA particles by a factor of 2 as compared to that observed in our earlier study of CMG-forked DNA in the absence of Mcm10 (Georgescu et al., 2017). We went on to determine a 3.9-Å cryo-EM structure (**Fig. 1a-d, Supplementary Fig. 1-5, Table 1**). To our surprise, no Mcm10 density was observed in the cryo-EM 3D map, suggesting that Mcm10 has multiple conformations or binds at multiple sites and becomes averaged out in the 3D reconstruction process. Because Mcm10 density is not observed, we will therefore refer to our 3D reconstructed structure as CMG-forked DNA in this manuscript, although we presume Mcm10 may be present. In the new structure, the parental dsDNA enters CMG with an angle essentially the same as in the previous lower resolution structure in the absence of Mcm10 (**Fig. 1c**). We suggest that Mcm10 binding does not alter the in-line configuration of the parental dsDNA and the unwound leading ssDNA. Therefore, the DNA configuration in the eukaryotic replisome is likely different from the T7 replisome. This different orientation, in-line vs perpendicular, appears important to ease of unwinding (Ribeck et al., 2010) and may contribute to the 10-100 fold different velocities of unwinding between eukaryotic and prokaryotic replicative helicases.

**Table 1.**
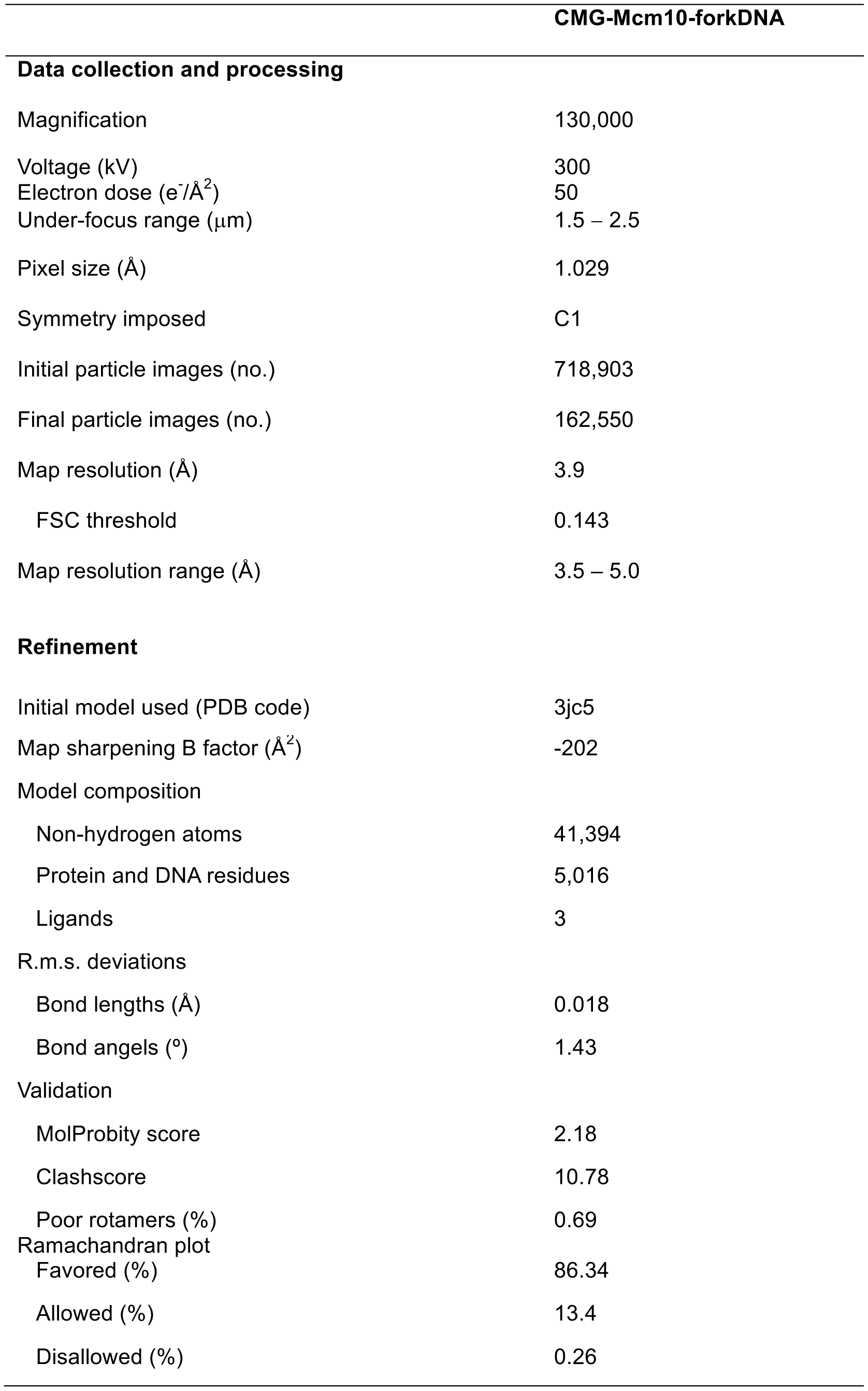
Cryo-EM 3D reconstruction and refinement of the CMG-forked DNA complex.

**Figure 1.**
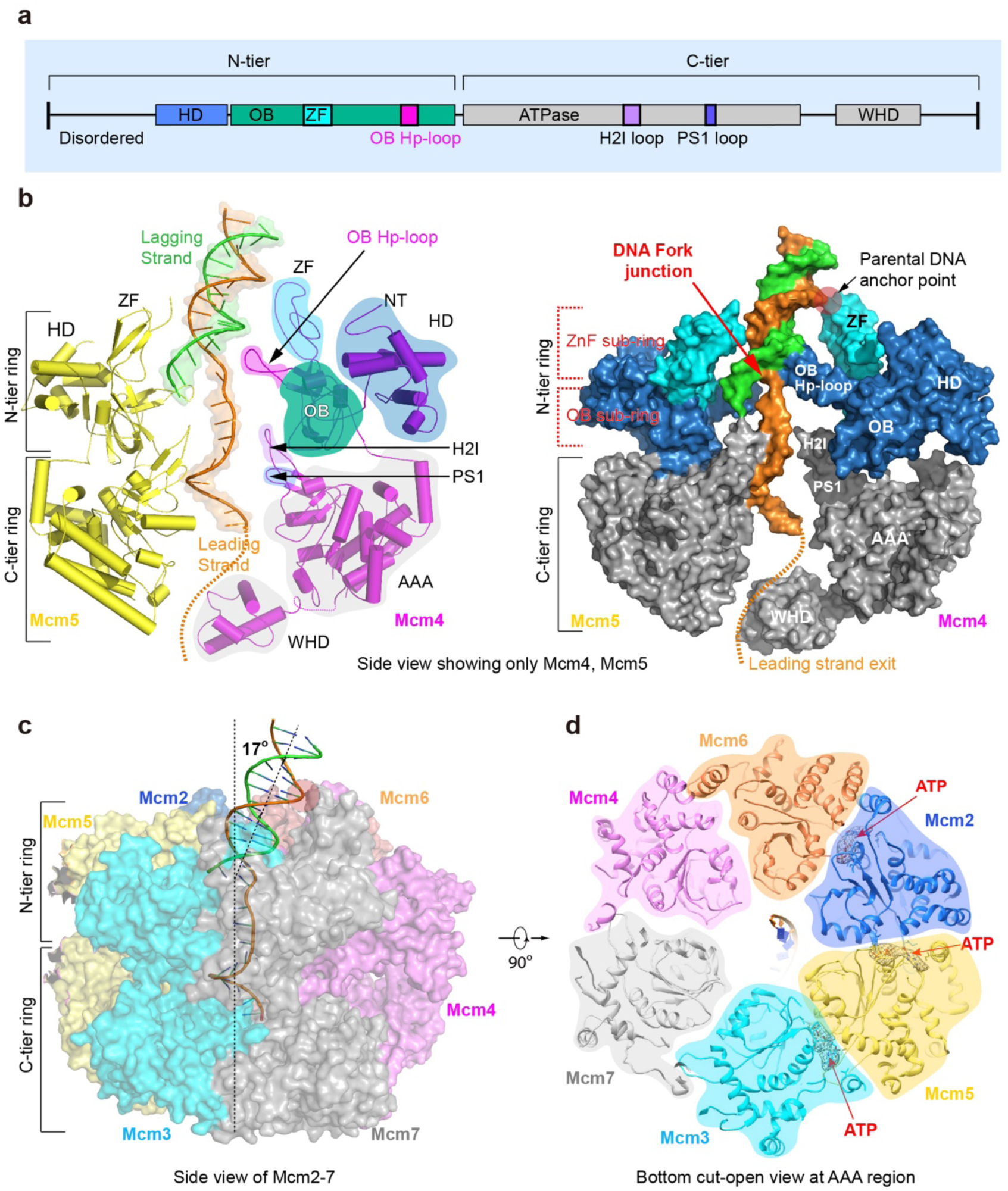
The parental dsDNA enters the zing finger region at the N-tier ring of CMG and is in-line with the unwound leading ssDNA. (**a**) A sketch of the general domain architecture of a typical Mcm protein, highlighting the hairpin loop in the OB domain (OB Hp-loop), and the H2I and PS1 loops in the AAA+ domain. HD: N-terminal helical domain, ZF: Zinc finger domain that is embedded in the OB domain, WHD: C-terminal winged helix domain. (**b**) Two side views of CMG-forked DNA complex in cartoon (left) and surface (right) views. Only Mcm4 and Mcm5 are shown to provide an unobstructed view of the forked DNA and the major structural features of a typical Mcm protein, as labeled in Mcm4. (**c**) A side view of the full CMG-forked DNA complex in transparent surface rendering, with DNA in cartoon. The two dashed lines show that the parental dsDNA is nearly in-line with the unwound leading ssDNA. (**d**) A bottom view of Mcm2-7 that is cut open at the AAA+ motor ring. Cryo-EM densities for the three ATP molecules are shown as gray mesh, superimposed with atomic model of ATP in sticks.

The improved structure contained three ATP molecules bound to Mcm2, Mcm3, and Mcm5, the right half of the Mcm ring embraced by Cdc45 and GINs (**Fig. 1d**). This is consistent with the three-nucleotide binding in the earlier study, although the identities of the nucleotides were undetermined in the previous low-resolution work (Georgescu et al., 2017). The structure clearly shows that only four subunits interact with the leading strand ssDNA in the C-tier ATPase motor ring: Mcm3, Mcm5, Mcm2, and Mcm6, while the Mcm4 and Mcm7 subunits do not bind DNA in the ATPase region (to be described below). Therefore, the overall structure underscores the asymmetry in the translocating motor ring in terms of ATP binding as well as in DNA binding. This property of the eukaryotic helicase seems to be different from the archaeal MCM hexamer as observed in the recent crystal structure of a mutant SsoMCM in which the linker between the N-tier and C-tier rings are shortened but the helicase is nevertheless active (Meagher et al., 2019). In such modified archaeal MCM hexamer structure, three Mcm proteins bound to ADP and three remaining subunits bound to ADP-BeF_3_, but all six proteins made contacts with the 12-base ssDNA in the central channel.

### The parental dsDNA reaches the boundary between the ZF sub-ring and OB sub-ring

CMG, in the context of a full replisome in Xenopus extracts, is demonstrated to act by a steric exclusion process, in which the DNA is split before entering the helicase (Fu et al., 2011). Thus, it was somewhat unexpected that dsDNA enters CMG a short distance at the Zinc finger (ZF) region, as observed in the CMG-forked DNA structure (Georgescu et al., 2017) as well as in a subsequent structure of CMG−Pol ε−ATPγS on the forked DNA (Goswami et al., 2018). The ZF domains of CMG project from the extreme N-face of CMG, even though the sequence of a helical domain (HD) or A-domain in the NTD of MCM subunits comes earlier in the primary sequence (**Fig. 1a**), the ZF and an OB fold are intertwined in the primary sequence and come directly after the A-region (**Fig. 1b**) (Miller and Enemark, 2015). In the current 3.9-Å structure, the ZFs project from the N-tier of Mcm2-7 and encircle and contact the dsDNA (**Fig. 1b**). Therefore, both the future leading and lagging strand DNA, while still parental duplex, are in fact contacted by CMG before their separation into single strand DNAs. However, the binding is not extensive, and only proceeds to the floor of the ZFs, after which the stands are separated. This separation point will be examined further in the next section. Physical interaction of CMG in the dsDNA region may explain why bulky groups that are held close, within 6-10 Å, to the duplex inhibited helicase activity (Langston et al., 2017).

### The OB hairpin (Hp) loops form a dam-like barrier that blocks the lagging strand

At the improved resolution of 3.9 Å, new DNA contacts in the N-tier ring reveal how the dsDNA is split apart during the ATP-driven translocation by the C-tier motors. The region in the Mcm6 OB domain from residue 403 to 453 undergoes a dramatic conformational change as compared to the apo CMG (**Fig. 2a**). In the absence of DNA these Mcm6 OB loop residues adopt a structure with two β-strands that is a substantial distance from the central channel of CMG. Upon helicase activity on forked DNA with ATP in the current structure, this Mcm6 OB loop region changes to an *α*-helix that projects the hairpin (Hp) loop 12 Å towards the central channel to interact with the DNA forked junction region. This Mcm6 OB Hp loop, together with OB Hp loops of Mcm4 and Mcm7, form a continuous and slanted barrier below the lagging strand (**Fig. 2b**). Thus, as the leading strand is pulled from below in the C-tier motor, the lagging strand at the fork is blocked by this barrier and can no longer move downwards along with the leading strand 5’ flap. Instead, the lagging strand flap is diverted sideways, and it is in this manner that the 5’ flapped duplex DNA is unwound and separated to the outside of CMG for steric exclusion unwinding (**Fig. 2c**).

**Figure 2.**
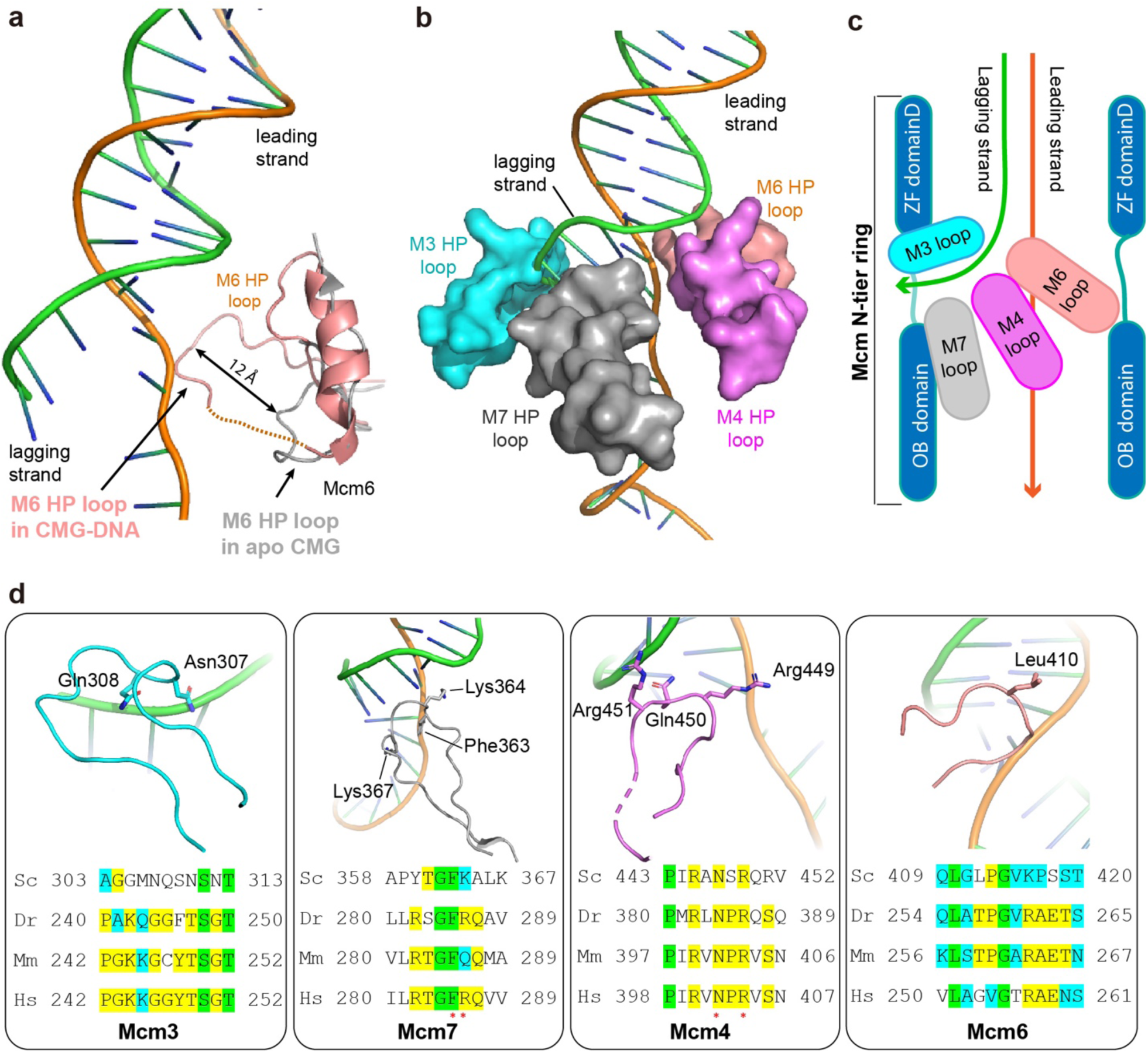
Interactions between four OB Hp-loops and DNA at the fork junction. **(a)** The Mcm6 OB hairpin loop undergoes a major conformation change upon binding to DNA. In the absence of DNA (grey) this region is largely disordered. In the presence of DNA (wheat), this region forms an *α*-helix that projects the OB Hp loop ∼12 Å towards the center to interact with the DNA fork. **(b)** The four OB Hp loops surrounding the forked DNA. **(c)** A sketch showing that the three OB Hp loops (of Mcm4, Mcm4, and Mcm7) spiral below thus blocking the lagging strand, while one OB Hp loop (of Mcm3) is above the lagging strand and may prevent the lagging strand from backing out from the top entry of the Mcm2-7 channel. (**d**) Zoomed views of the four fork-interacting OB Hp loops with key residues shown in sticks. Below each panel is shown a sequence alignment of the OB Hp loop from Sc – *Saccharomyces cerevisiae*, Dr -*Danio rerio*, Mm – *Mus musculus*, and Hs – *Homo sapiens*. Red asterisks below the sequence mark invariant residues.

Interestingly, unlike the OB Hp loops of Mcm6, Mcm4, Mcm7 that are below the lagging strand, the OB Hp loop of Mcm3 contacts the unwound lagging strand from above (**Fig. 2b-d**). This strategic position prevents the lagging strand from making a U-turn to back out of the central chamber from the top entry. Therefore, the four OB Hp loops together appear to form a diversion tunnel, with three OB Hp loops of Mcm7, 4, and 6 forming the lower wall, and the single OB Hp loop of Mcm3 forming the upper wall, thereby guiding the lagging strand towards the side to exit the helicase.

Comparison of sequences of the four OB Hp loops of Mcm3, Mcm4, Mcm6, and Mcm7 indicate some degree of conservation among eukaryotes (**Fig. 2d**). The Mcm7 OB Hp loop projects two lysine residues (Lys-364 and Lys-367) and one conserved and bulky phenylalanine (Phe-363) towards the fork junction. The Mcm4 OB Hp loop projects two conserved arginine residues (Arg-449 and Arg-451) and one conserved polar residue (Gln-450) towards the junction. However, the interactions between the OB Hp loops of Mcm3 and Mcm6 are less conserved, involving polar and small hydrophobic residues that project toward the junction.

### A possible exit port for the lagging strand DNA

In the current structure, the density of the lagging strand past the fork junction is lost at the base between the ZF domains of Mcm3 and Mcm5. Several structural features suggest that this region forms a possible exit path for the unwound lagging strand. Firstly, the Mcm3 subunit lacks a Zn atom, a conserved modification of Mcm3 subunits in eukaryotes (Li and O’Donnell, 2018; O’Donnell and Li, 2018), and likely explains why the ZF domain of Mcm3 appears collapsed in the structure compared to those of other Mcm subunits that contain a Zn atom (**Fig. 3a-b**). Secondly, a sizable gap between ZF domains of Mcm3 and Mcm5 – required for the potential passage of the lagging strand – is maintained by the N-terminal loop (aa 1-14) of the neighboring Mcm7, which is well ordered and reaches over to interact with the Mcm3 ZF domain. The N-terminal 100-200 residues of an Mcm protein is typically disordered (**Fig. 1a**); thus it is unexpected that the extreme N-terminus of Mcm7 is well ordered and reaches over to interact with, and stabilize the Mcm3 ZF domain (**Fig. 3a,b**). Thirdly, the floor of the gap between the ZF domains of Mcm3 and Mcm5, formed by an extended insertion hairpin loop of the Mcm3 ZF domain is well conserved, as highlighted by blue and red boxes in **Fig. 3c**. Furthermore, on the left side of the gap, the Mcm5 ZF domain projects two conserved arginine residues (Arg-184 and Arg-187) towards the gap to possibly interact with the lagging strand (**Fig. 3a, c**). We have tried site specific modification of important residues, such as the ATP site Walker A motifs of recombinant Mcm1-7 genes, but considering that Crisper technology will modify both the endogenous copy and the recombinant copy, we obtained negative results (yet we see that Crisper worked from the silent mutations we placed in the gRNA and confirmed they were present in product cells). Thus essential changes are not able to be analyzed without adopting a heterogeneous expression method (i.e. baculovirus). Regardless, deletion and substitution (with Gly-Ala) of the OB loops could not be obtained consistent with an important function of the floor region. Taken together, the lagging strand is likely shunted out of CMG after entering the ZF “tower” over the collapsed floor between the ZF domains of Mcm3 and Mcm5, prior to entry into the bona fide central channel of CMG.

**Figure 3.**
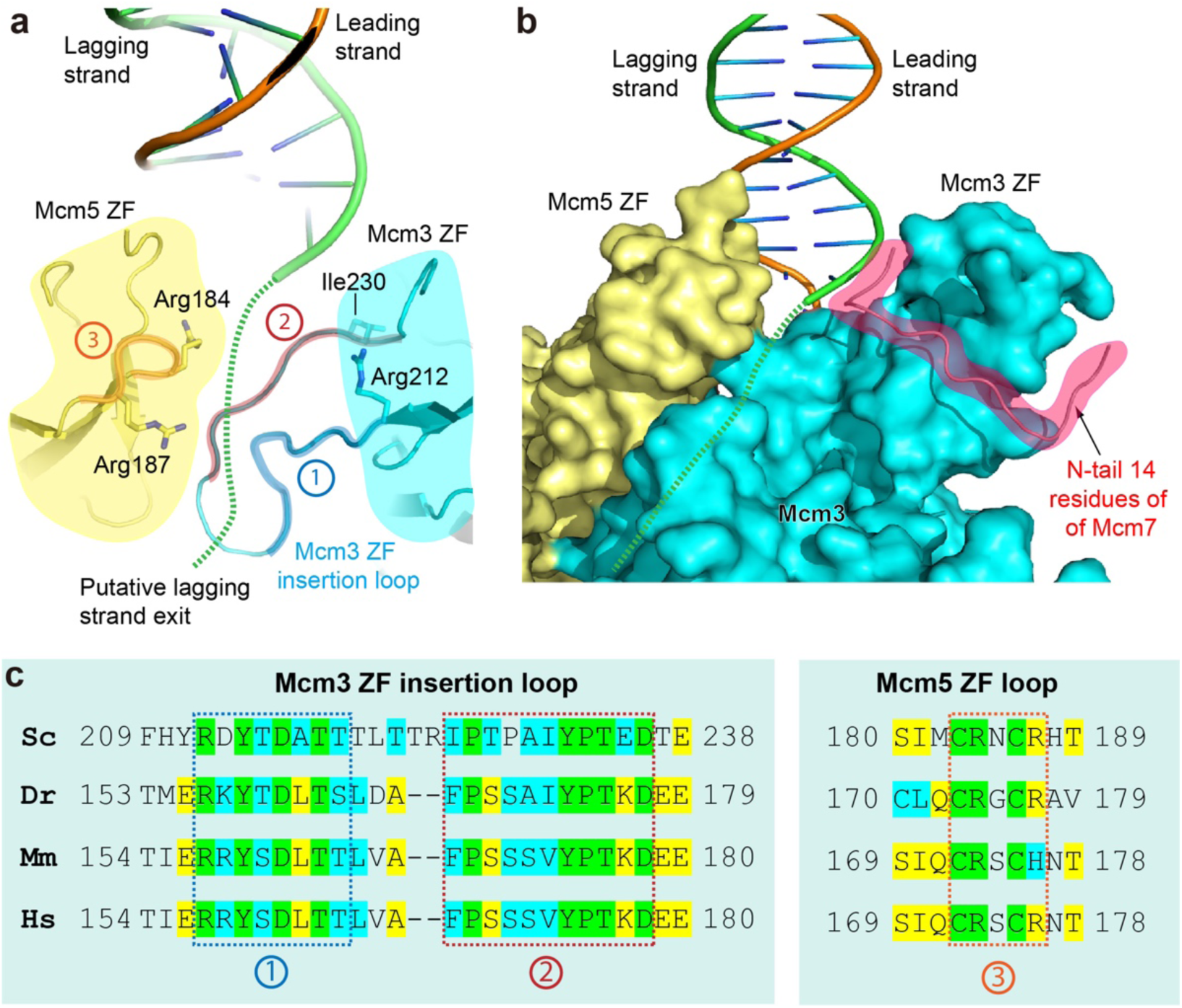
A putative exit groove of lagging strand DNA between ZF domains of Mcm3 and 5. (**a**) Close up of the DNA fork within the ZF regions, with the position of cryo-EM density loss of the lagging strand (dashed green curves) positioned toward the viewer. This density loss occurs at a groove between ZF domains of Mcm3 and 5. The floor of the groove is lined by a long hairpin loop (labeled by 1 and 2 inside red circles) of Mcm3 ZF domain, which has the unique and conserved feature among eukaryotic CMGs of lacking a Zn atom. A loop in Mcm5 ZF domain (labeled by “3” in a red circle) forms the left wall of the groove and projects two Arg residues towards the putative lagging strand groove. (**b**) The gap between ZF domains of Mcm3 and 5 is sustained by the ordered N-terminal 12-residue peptide of the neighboring Mcm7, shown in cartoon and highlighted in red shade. This peptide forms the right wall of the groove. (**c**) Sequence alignment of the groove-forming loops of the ZF domains of Mcm3 and Mcm5. Sc: *Saccharomyces cerevisiae*, Dr: *Danio rerio*, Mm: *Mus musculus*, and Hs: *Homo sapiens*. Green denotes invariant residues, yellow denotes 3 out of 4 residues are conserved. Blue indicates residues having similar properties.

### The unwound leading strand is stretched straight at the fork junction

Under relaxed and normal conditions, the conformation of ssDNA is more condensed than dsDNA because it has many more degrees of freedom and can take on various structures that dsDNA does not have available (Bustamante et al., 1994). This is clearly not observed for the leading ssDNA within the N-region of CMG that is nearly linear (**Fig. 1b, Fig. 4a**). The motors of CMG are in the C-tier, below the N-tier during duplex DNA unwinding. Thus, one can deduce that that the CMG motors in the C-tier pull the leading strand DNA such that the parental lagging strand is forced against the slanted OB loop barrier leading to the diversion of the lagging strand towards the side port as discussed above. Importantly, there is little interaction between the N-tier linear segment of the leading strand and the helicase. Thus, the nearly linear conformation of the leading ssDNA in the N-tier channel can only be explained by the pulling force of the motors in the C-tier below. This structural feature is further evidence for the steric exclusion model, in which DNA is separated at the top N-tier of the helicase – between the ZF ring and OB ring – and not in the middle between the N- and C-tiers.

**Figure 4.**
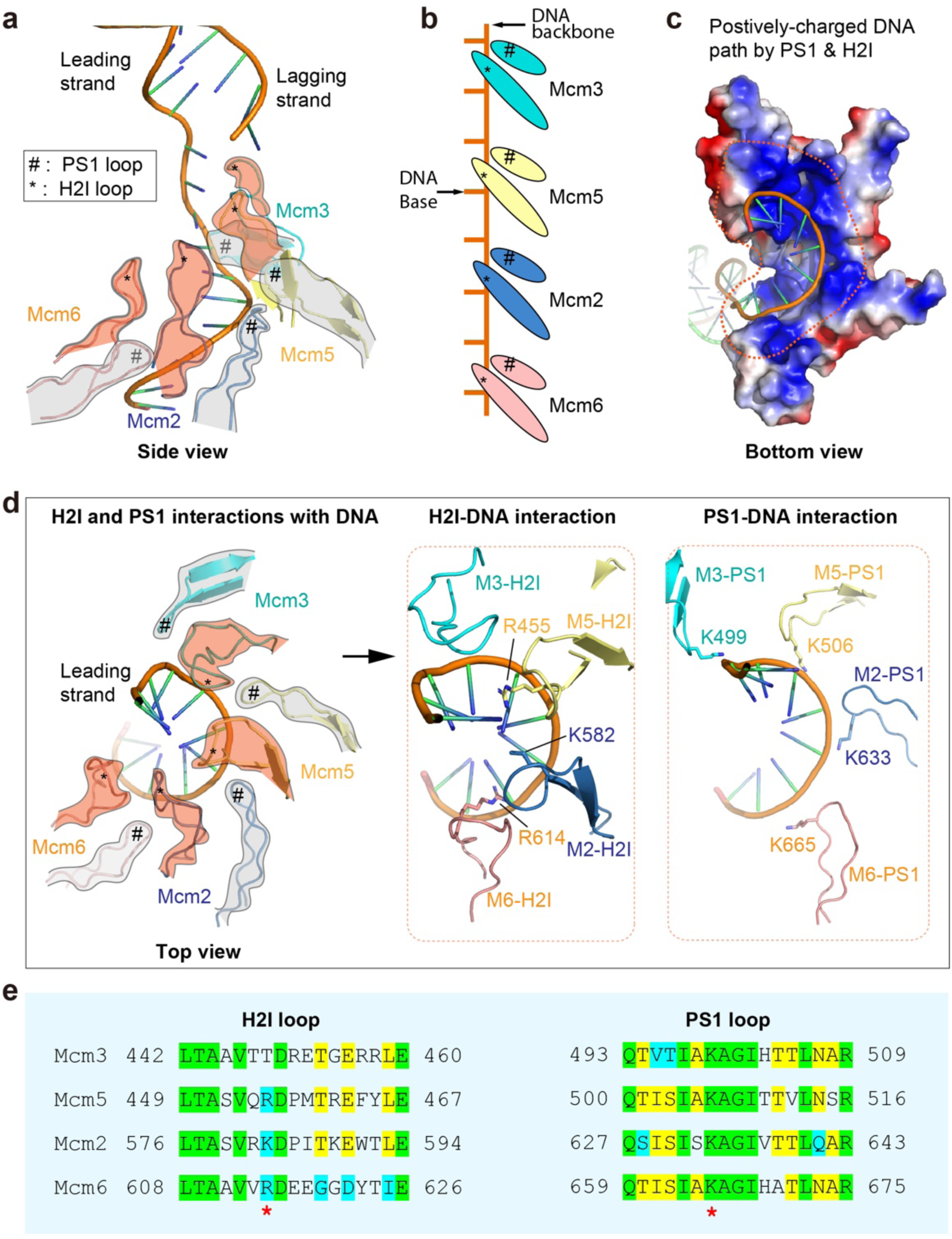
The leading ssDNA inside the C-tier motor ring of CMG helicase. (**a**) A side view of the DNA in the CMG chamber. The DNA-translocating PS1 and H2I loops of only four Mcm subunits (Mcm6, Mcm2, Mcm5, and Mcm3) spiral around and make contact with the leading ssDNA. (**b**) An illustration showing that among the four Mcm subunits engaging the DNA, each PS1 binds two phosphates PS1 loop along with an H2I loop binding over a base. The DNA-translocating loops rise two bases per Mcm subunit. (**c**) A bottom view showing the positively charged spiral path of the leading ssDNA formed by the PS1 and H2I loops of Mcm2, Mcm3, Mcm5, and Mcm6. (**d**) A top view of the PS1 and H2I loops in Mcm 3, 5, 2 and 6 engaging the spiral leading ssDNA. The middle and right panels show the DNA-interacting positively charged residues of H2I (middle) and PS1 (right). (**e**) Sequence alignment of the H2I and PS1 loops among the four DNA-interacting Mcm subunits. Red asterisks mark the conserved and positively charged residues.

### Binding of the leading ssDNA to the C-tier motor domains

The AAA+ motif of the CMG ATP site (SF6 helicase) contains loops that bind to DNA, referred to as PS1 and H2I (Erzberger and Berger, 2006). The PS1 loops are thought to be the main loops that act in the ATP-driven translocation complex, because the related eukaryotic viral BPV E1 and SV40 T-antigen, helicases in the SF4 class, are AAA+ helicases that contain PS1 loops but lack H2I loops (Erzberger and Berger, 2006). Although each of the six MCM proteins has a PS1, only four PS1 loops interact with the leading ssDNA in our structure (i.e. the PS1 loops of Mcm3, Mcm5, Mcm2, and Mcm6 (**Fig. 4a-e**)). Each of the four PS1 loops binds two phosphodiester backbone links (**Fig. 4b**). This two-phosphodiester-per-PS1 binding mode also applies to the bacterial DnaB and phage T7 helicase, as revealed by recent high-resolution structures (Gao et al., 2019; Kose et al., 2019). This has important implications in generalizing helicase activity from bacteria to eukaryotes as explored further in the Discussion. In contrast, the H2I element, unique to the SF6 class to which CMG belongs, is observed in the current structure to interact with a nucleotide base located between the two phosphodiester bonds bound by the PS1 loop, and not the phosphate backbone (**Fig. 4d**). The four H2I loops join the 4 PS1 loops to form a highly positively charge and well-conserved spiral path for the DNA spiral in the motor region (**Fig. 4c, e**). In our CMG-forked DNA structure, the manner in which the four H2I loops interact with nucleotide bases is essentially the same, and implies that the H2I interaction to DNA is an important element of DNA translocation.

## DISCUSSION

The improved resolution of 3.9 Å has provided an unprecedented visualization of a fork junction in a replicative DNA helicase. The structure shows that the parental duplex enters the ZF sub-ring, and the strand separation occurs at the bottom of the ZF sub-ring and the beginning of the OB sub-ring. The unwound leading ssDNA traverses the OB sub-ring almost linearly in the N-tier but spirals through the C-tier motor ring before emerging from the C-face of the CMG helicase. Our earlier studies revealed that bulky lagging strand blocks inhibited the CMG helicase, consistent with structural studies showing that dsDNA enters the helicase (Langston et al., 2017). However, use of >40-Å multi-PEG spacer linkers to tether bulky groups to the lagging strand did not inhibit CMG (Kose et al., 2019). We presume that these long linkers enable the lagging strand base to reach the fork junction inside CMG and can be unwound while keeping the steric block outside of the surface of the CMG helicase. The height of the ZF collar is about 30 Å, making such scenario possible.

### In-line geometry of parental dsDNA and unwound leading ssDNA

The dsDNA entering CMG in the presence of ATP is similar to the recent CMG-Pol ε-forked DNA structure using ATPγS (Goswami et al., 2018). Hence, the evidence would suggest that DNA enters CMG in-line, with the parental duplex being held by the ZF collar, unobstructed until reaching to the OB sub-ring. The ZF collar in the eukaryotic CMG apparently sets the approaching angle of the parent DNA and may further protect the fork from other helicases like Pif1 or Rrm3 during normal replication.

The duplex is oriented at a right angle to the ssDNA entering the helicase chamber In the T7 gp4 replisome (Gao et al., 2019). The gp4 helicase has a very short N-tier, similar to other bacterial and some viral hexameric helicases (O’Donnell and Li, 2018). Thus, it would appear that the ability to bind dsDNA is not a general element of helicase action. Interestingly, it is proposed that the perpendicular arrangement of ds and ssDNA in the T7 helicase-forked DNA structure is important for rapid unwinding (Gao et al., 2019). This statement is reasonable because bacterial forks travel 1-2 orders of magnitude faster than eukaryotic forks, and the geometry of the DNA is shown to affect the force needed to unzip it (Ribeck et al., 2010). Perhaps the slow speed of eukaryotic forks eliminated the selection pressure to maintain a perpendicular geometry of the dsDNA-ssDNA junction and explaining the in-line geometry observed in our and others study of CMG-DNA (Douglas et al., 2018; Georgescu et al., 2017; Goswami et al., 2018).

### A “diversion tunnel” model of steric exclusion DNA unwinding

The term “steric exclusion” implies there is no specific unwinding element, other than the fact that one strand of the duplex can not fit in (i.e. is sterically excluded from) the central channel of the helicase. However, we have shown previously that Saccharromyces cerevisiae CMG can traverse dsDNA (Wasserman et al., 2019). The current study clarifies the detailed interactions between the DNA fork junction and CMG, leading us to propose the following “diversion-tunnel” model for steric exclusion DNA unwinding by CMG (**Fig. 5**). In this model, the three OB loops of Mcm7, 4, and 6 form a dam just below the lagging strand at the fork nexus, and the OB loop of Mcm3 located above the lagging strand, together with those of Mcm4, 7, and 6, form the diversion tunnel via which the lagging strand is guided sideways, and may exit the CMG chamber via a gap between the ZF domains of Mcm3 and Mcm5. The driving force for the duplex unwinding is then the pulling force of the C-tier motors on the leading strand DNA from below.

**Figure 5.**
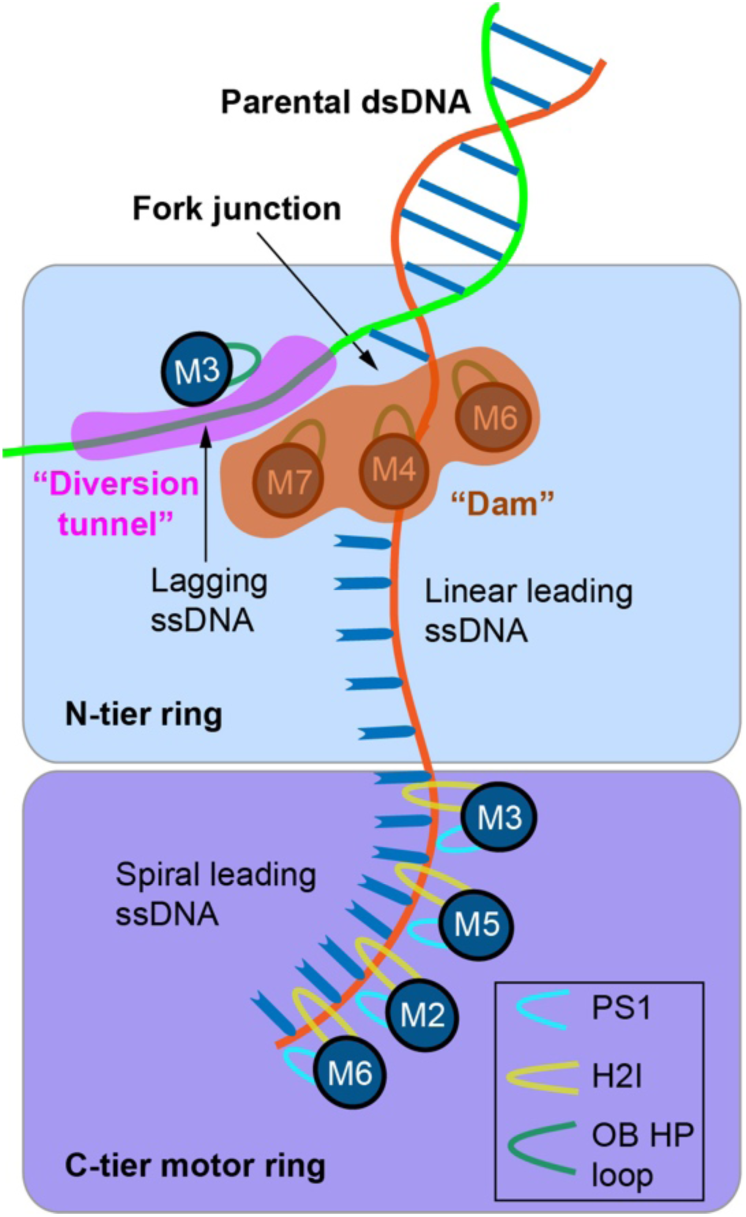
Model for DNA unwinding derived from this study. OB Hp loops in the NTDs of Mcms7, 4 nd 6 form a block, or dam for incoming 5’ tailed DNA, and the OB Hp loop in the NTD of Mcm3 forms the top of a diversion tunnel, or channel for the displaced lagging 5’ strand. The PS1 and H2I loops in the C-tier motor domain pull the DNA through the diversion tunnel in the N-tier for the steric exclusion process of DNA unwinding. See text for details.

Our model is different from the widely anticipated “separation pin” model of steric exclusion, in which a specific structural element is responsible for DNA unwinding, which would in turn predict an absolutely conserved structural element or amino acid sequence among all eukaryotic helicases. Such an absolutely conserved sequence has never been identified and may not exist. In our “diversion-tunnel” model, the structural elements – the OB loops that actually form the dam and the tunnels – are largely structural and require neither specific amino acid sequences nor specific interactions between the forked DNA and the unwinding apparatus.

### The PS1 and H2I loop motifs

We note that only four of the six subunits of CMG bind ssDNA in the six AAA+ modules, similar to our lower resolution study (Georgescu et al., 2017) and to the CMG-Pol ε-fork-ATPγS study (Goswami et al., 2018). In contrast, the homo-hexameric replicative helicases appear to bind ssDNA with all six of the AAA+ domains (Li and O’Donnell, 2018; Meagher et al., 2019). Why only four, but not all six, PS1 loops in CMG simultaneously pull on the leading ssDNA is currently not understood. It is possible that we have captured just a single pose and the two unbound PS1 loops will engage the leading ssDNA during a full translocation cycle. This possibility is supported by the fact that the DNA-interacting PS1 loops are from Mcm2, Mcm3, Mcm5, and Mcm6 in the ATP-bound structure, but are from Mcm4, Mcm6, and Mcm7 in the ATPγS-bound structure (Abid Ali et al., 2016).

The two-phosphates-per-PS1 DNA binding mode observed in CMG is shared with the *E. coli* DnaB helicase and the phage T7 helicase (Itsathitphaisarn et al., 2012; Gao et al., 2019), and this DNA binding mode is further reported in the *E. coli* clamp loader AAA+ ring ATPases (Simonetta et al., 2009). Thus, the binding of two phosphodiester bonds per subunit in a AAA+ motor may often be used by replicative helicases and possibly other DNA helicases and translocases. However, helicases that bind only one phosphodiester link per translocation loop have also been described, such as the *E. coli* Rho factor and Bovine Papilloma Virus E1 helicase (Enemark and Joshua-Tor, 2006; Thomsen and Berger, 2009). It is currently unclear why some helicases translocate on DNA one base at a time, while others like CMG translocate two bases per step.

The function of H2I is less well understood. Although the H2I motif is not present in SF4 AAA+ helicases, this motif is required for activity in the archaeal MCM helicase (Jenkinson and Chong, 2006). The H2I loops clamp down on the leading strand in the crystal structure of archaeal MCM (Miller et al., 2014). In our CMG structure, each of the four H2I loops of Mcm3, Mcm5, Mcm2, and Mcm6 contacts a nucleotide base, in sync with the four PS1 loops of the same Mcm subunits contacting the phosphodiester backbone, as if the H2I loops play a secondary yet essential role in support of DNA translocation by the PS1 loops.

### Does CMG function by staircasing?

It is generally accepted that the C-tier motors move like a staircase in homo-hexameric archaeal helicases (Enemark and Joshua-Tor, 2006, 2008; Lyubimov et al., 2011; Thomsen and Berger, 2009). How C-tier motors translocate DNA in the eukaryotic CMG is largely unknown, but mechanistic differences between the archaeal and eukaryotic helicases may be expected, because each Mcm subunit is a distinct protein in CMG, and In vitro mutational studies suggest that only two sites are essential for helicase activity: the site located between Mcm2 and Mcm5, in which subunit Mcm5 binds the ATP, and the site between Mcm3 and Mcm5 in which Mcm3 binds the ATP (Ilves et al., 2010). However all the Mcm subunit ATP Walker A sites appear required for robust cell viability (Coster et al., 2014; Kang et al., 2014). It is possible that the ATPase activity of certain Mcm proteins may be essential for their loading onto DNA by origin recognition complex, in concert with Cdc6 and Cdt1 or during origin activation where the inactive double Mcm2-7 hexamers are converted into two active CMG helicases (Riera A et al., 2017; Douglas et al., 2018). Alternatively, in the context of the moving replisome with Mcm10, DNA polymerases and additional subunits present, all or most Mcm ATP sites might be utilized, and thus a staircasing process of unwinding can not be ruled out.

We and others previously suggested at least two non-staircasing motions that could move DNA for the purpose of CMG translocation: either the C-tier and N-tier move relative to one another (Abid Ali et al., 2016; Li and O’Donnell, 2018; Yuan et al., 2016), or two C-tier motor subunits, Mcm2 and 5, move relative to one another in an inch-worming mechanism (Li and O’Donnell; Zhai et al, 2017). However, the C-tier/N-tier movement is thus far only observed in the apo CMG (Yuan et al., 2016) and not yet observed in CMG bound to DNA such as in the current structure, suggesting the pumpjack is less likely to be operational. Furthermore, a recent study of CMG-Pol ε bound to ATPγS at a DNA fork suggests that the C-tier AAA+ motor domains are planar rather than spiral, closing the gap at the motor domains of Mcm2 and 5 (Georgescu et al., 2017; Goswami et al., 2018). The current CMG-forked DNA structure in the presence of ATP also shows a compact planar form at the C-tier, and thus we conclude that Pol ε is not required for the planar conformation observed in the presence of Pol ε (Georgescu et al., 2017; Goswami et al., 2018). Assuming that movement of the Mcm2 and 5 interface is incompatible with (or without) Pol ε binding, inch-worming translocation by movements between Mcm2 and Mcm5 may also be inoperable. Given the cell viability studies indicating all ATP sites play important roles in CMG function (Coster et al., 2014; Kang e. al., 2014), we don’t exclude the rotational staircasing model. Indeed one cryoEM structure of fly CMG-DNA-ATPγS shows leading ssDNA interacts with Mcms 7,4,6 (Abid Ali et al., 2016), while the leading ssDNA binds Mcms 6,2,5,3 in the current report and in other studies (Georgescu et al., 2017; Goswami et al., 2018). Thus, it is possible all subunits bind ssDNA during the full catalytic cycle of CMG translocation, and further studies are clearly needed to address this. Moreover, It is interesting to note that the BPV E1 helicase and Rho helicase are planar, and only the DNA binding loops move in a spiral for staircasing (Enemark and Joshua-Tor, 2006, 2008; Lyubimov et al., 2011; Thomsen and Berger, 2009), distinct from E. coli DnaB (Itsathitphaisarn et al., 2012) and T7 gp4 (Gao et al., 2019) that move entire ATP binding domains during staircasing. Hence, if CMG acts by a staircase mechanism, it may do so as a planar disk, like E1 and Rho, with only the DNA binding loops performing the translocation movements rather than whole domain movements.

In our new structure, only three nucleotide pockets are occupied by ATP, with a forth site partily occupied by an ATP or ADP, and the remaining two sites are empty. In the sequential staircasing mechanism, all six ATP sites are obligated to cycle through ATP, ADP, and apo (empty) configurations, but only one site is expected to be empty at any given time in the hexameric motor ring. In this scenario, the observed two empty sites may be explained by the loss of one nucleotide (likely an ADP) during cryo-EM sample preparation, as the buffer solution contained only ATP but no ADP. However, it is also possible that the ATPase activity of only some Mcm proteins function in DNA unwinding. In this case, the two empty sites we have observed may be non functional for the helicase activity. If true, this would imply a stochastic unwinding mechanism such as the lazy Brownian ratchet mechanism, proposed recently based on a single molecule study of the Drosophila CMG (Burnham et al., 2019). The stochastic model of DNA unwinding has its root in the probabilistic ATP hydrolysis sequence first discovered in the peptide unfoldase ClpX (Martin, et al., 2005). To summarize, the current report sheds light on a dam-and-diversion process explaining how steric exclusion may function for CMG, and further studies will be needed to understand the translocation mechanism of CMG.

## Acknowledgement

Cryo-EM data were collected at the David Van Andel Advanced Cryo-Electron Microscopy Suite at the Van Andel Research Institute. We thank G. Zhao and X. Meng for help with data collection. This study was supported by the US National Institutes of Health grants GM131754 (to H.L.) and GM115809 (to M.E.O), and the Howard Hughes Medical Institute (M.E.O).

## Author contributions

Z.Y. R.G, H.L., and M.E.O conceived and designed experiments. R.G. and D.Z. purified protein, reconstituted CMGM and prepared samples of CMGM-forked DNA. Z.Y. performed the EM experiments. Z.Y. and L.B. performed image processing and atomic modelling. Z.Y., R.G., L.B., H.L, and. M.E.O analyzed the data. Z.Y., H.L., and M.E.O wrote the manuscript.

## Accession codes

The 3D cryo-EM maps of CMG-forked DNA at 3.9 Å resolution has been deposited in the Electron Microscopy Data Bank under accession code EMD-20607. The corresponding atomic model has been deposited in the Protein Data Bank under accession code PDB-6U0M.

## Literature cited

Abid Ali, F., Douglas, M.E., Locke, J., Pye, V.E., Nans, A., Diffley, J.F.X., and Costa, A. (2017). Cryo-EM structure of a licensed DNA replication origin. Nat Commun 8, 2241.

Abid Ali, F., Renault, L., Gannon, J., Gahlon, H.L., Kotecha, A., Zhou, J.C., Rueda, D., and Costa, A. (2016). Cryo-EM structures of the eukaryotic replicative helicase bound to a translocation substrate. Nat Commun 7, 10708.

Burnham DR, Kose HB, Hoyle RB, and Yardimci H. (2019) The mechanism of DNA unwinding by the eukaryotic replicative helicase. Nat Commun. 10, 2159.

Bustamante, C., Marko, J.F., Siggia, E.D., and Smith, S. (1994). Entropic elasticity of lambda-phage DNA. Science 265, 1599–1600.

Costa, A., Renault, L., Swuec, P., Petojevic, T., Pesavento, J., Ilves, I., MacLellan-Gibson, K., Fleck, R.A., Botchan, M.R., and Berger, J.M. (2014). DNA binding polarity, dimerization, and ATPase ring remodeling in the CMG helicase of the eukaryotic replisome. Elife, e03273.

Coster, G., Frigola, J., Beuron, F., Morris, E.P., and Diffley, J.F. (2014). Origin licensing requires ATP binding and hydrolysis by the MCM replicative helicase. Mol Cell 55, 666–677.

Davey, M.J., Indiani, C., and O’Donnell, M. (2003). Reconstitution of the Mcm2-7p heterohexamer, subunit arrangement, and ATP site architecture. J Biol Chem 278, 4491–4499.

Douglas, M.E., Ali, F.A., Costa, A., and Diffley, J.F.X. (2018). The mechanism of eukaryotic CMG helicase activation. Nature 555, 265–268.

Douglas, M.E., and Diffley, J.F. (2016). Recruitment of Mcm10 to Sites of Replication Initiation Requires Direct Binding to the Minichromosome Maintenance (MCM) Complex. J Biol Chem 291, 5879–5888.

Enemark, E.J., and Joshua-Tor, L. (2006). Mechanism of DNA translocation in a replicative hexameric helicase. Nature 442, 270–275.

Enemark, E.J., and Joshua-Tor, L. (2008). On helicases and other motor proteins. Curr Opin Struct Biol 18, 243–257.

Erzberger, J.P., and Berger, J.M. (2006). Evolutionary relationships and structural mechanisms of AAA+ proteins. Annu Rev Biophys Biomol Struct 35, 93–114.

Fu, Y.V., Yardimci, H., Long, D.T., Ho, T.V., Guainazzi, A., Bermudez, V.P., Hurwitz, J., van Oijen, A., Scharer, O.D., and Walter, J.C. (2011). Selective bypass of a lagging strand roadblock by the eukaryotic replicative DNA helicase. Cell 146, 931–941.

Gao, Y., Cui, Y., Fox, T., Lin, S., Wang, H., de Val, N., Zhou, Z.H., and Yang, W. (2019). Structures and operating principles of the replisome.Science 363.

Georgescu, R., Yuan, Z., Bai, L., de Luna Almeida Santos, R., Sun, J., Zhang, D., Yurieva, O., Li, H., and O’Donnell, M.E. (2017). Structure of eukaryotic CMG helicase at a replication fork and implications to replisome architecture and origin initiation. Proc Natl Acad Sci U S A 114, E697–E706.

Goswami, P., Abid Ali, F., Douglas, M.E., Locke, J., Purkiss, A., Janska, A., Eickhoff, P., Early, A., Nans, A., Cheung, A.M.C., et al. (2018). Structure of DNA-CMG-Pol epsilon elucidates the roles of the non-catalytic polymerase modules in the eukaryotic replisome. Nat Commun 9, 5061.

Ilves, I., Petojevic, T., Pesavento, J.J., and Botchan, M.R. (2010). Activation of the MCM2-7 helicase by association with Cdc45 and GINS proteins. Mol Cell 37, 247–258.

Itsathitphaisarn, O., Wing, R.A., Eliason, W.K., Wang, J., and Steitz, T.A. (2012). The hexameric helicase DnaB adopts a nonplanar conformation during translocation. Cell 151, 267–277.

Jenkinson, E.R., and Chong, J.P. (2006). Minichromosome maintenance helicase activity is controlled by N-and C-terminal motifs and requires the ATPase domain helix-2 insert. Proc Natl Acad Sci U S A 103, 7613–7618.

Kang, S., Warner, M.D., and Bell, S.P. (2014). Multiple functions for Mcm2-7 ATPase motifs during replication initiation. Mol Cell 55, 655–665.

Kose, H.B., Larsen, N.B., Duxin, J.P., and Yardimci, H. (2019). Dynamics of the Eukaryotic Replicative Helicase at Lagging-Strand Protein Barriers Support the Steric Exclusion Model. Cell Rep 26, 2113–2125 e2116.

Langston, L.D., Mayle, R., Schauer, G.D., Yurieva, O., Zhang, D., Yao, N.Y., Georgescu, R.E., and O’Donnell, M.E. (2017). Mcm10 promotes rapid isomerization of CMG-DNA for replisome bypass of lagging strand DNA blocks. eLife 6, e29118.

Li, H., and O’Donnell, M.E. (2018). The Eukaryotic CMG Helicase at the Replication Fork: Emerging Architecture Reveals an Unexpected Mechanism. Bioessays 40.

Looke, M., Maloney, M.F., and Bell, S.P. (2017). Mcm10 regulates DNA replication elongation by stimulating the CMG replicative helicase. Genes Dev 31, 291–305.

Lyubimov, A.Y., Strycharska, M., and Berger, J.M. (2011). The nuts and bolts of ring-translocase structure and mechanism. Curr Opin Struct Biol 21, 240–248.

Martin A, Baker TA, and Sauer RT. (2005) Rebuilt AAA + motors reveal operating principles for ATP-fueled machines. Nature. 437, 1115–20.

Mayle, R., Langston, L., Molloy, K.R., Zhang, D., Chait, B.T., and O’Donnell, M.E. (2019). Mcm10 has potent strand-annealing activity and limits translocase-mediated fork regression. Proc Natl Acad Sci U S A 116, 798–803.

Meagher, M., Epling, L.B., and Enemark, E.J. (2019). DNA translocation mechanism of the MCM complex and implications for replication initiation. Nat Commun 10, 3117.

Miller, J.M., Arachea, B.T., Epling, L.B., and Enemark, E.J. (2014). Analysis of the crystal structure of an active MCM hexamer. Elife 3, e03433.

Miller, J.M., and Enemark, E.J. (2015). Archaeal MCM Proteins as an Analog for the Eukaryotic Mcm2-7 Helicase to Reveal Essential Features of Structure and Function. Archaea 2015, 305497.

Moyer, S.E., Lewis, P.W., and Botchan, M.R. (2006). Isolation of the Cdc45/Mcm2-7/GINS (CMG) complex, a candidate for the eukaryotic DNA replication fork helicase. Proc Natl Acad Sci U S A 103, 10236–10241.

O’Donnell, M.E., and Li, H. (2018). The ring-shaped hexameric helicases that function at DNA replication forks. Nat Struct Mol Biol 25, 122–130.

Parker, M.W., Botchan, M.R., and Berger, J.M. (2017). Mechanisms and regulation of DNA replication initiation in eukaryotes. Crit Rev Biochem Mol Biol 52, 107–144.

Ribeck, N., Kaplan, D.L., Bruck, I., and Saleh, O.A. (2010). DnaB helicase activity is modulated by DNA geometry and force. Biophys J 99, 2170–2179.

Riera A, Barbon M, Noguchi Y, Reuter LM, Schneider S, and Speck C. (2017) From structure to mechanism-understanding initiation of DNA replication. Genes Dev. 31, 1073–1088.

Simonetta, K.R., Kazmirski, S.L., Goedken, E.R., Cantor, A.J., Kelch, B.A., McNally, R., Seyedin, S.N., Makino, D.L., O’Donnell, M., and Kuriyan, J. (2009). The mechanism of ATP-dependent primer-template recognition by a clamp loader complex. Cell 137, 659–671.

Skordalakes, E., and Berger, J.M. (2006). Structural insights into RNA-dependent ring closure and ATPase activation by the Rho termination factor. Cell 127, 553–564.

Sun J, Shi Y, Georgescu RE, Yuan Z, Chait BT, Li H, O’Donnell ME (2015). The architecture of a eukaryotic replisome. Nature Struct Mol Biol 22, 976–82.

Thomsen, N.D., and Berger, J.M. (2009). Running in reverse: the structural basis for translocation polarity in hexameric helicases. Cell 139, 523–534.

Wasserman, M.R., Schauer, G.D., O’Donnell, M.E., and Liu, S. (2019). Replication Fork Activation Is Enabled by a Single-Stranded DNA Gate in CMG Helicase. Cell 178, 600–611 e616.

Yuan, Z., Bai, L., Sun, J., Georgescu, R., Liu, J., O’Donnell, M.E., and Li, H. (2016). Structure of the eukaryotic replicative CMG helicase suggests a pumpjack motion for translocation. Nat Struct Mol Biol 23, 217–224.

Zhai Y, Cheng E, Wu H, Li N, Yung PY, Gao N, Tye BK. (2017) Open-ringed structure of the Cdt1-Mcm2-7 complex as a precursor of the MCM double hexamer. Nat Struct Mol Biol. 24, 300–308.

