## Supplemental Methods and Figures for "DNA unwinding mechanism of a eukaryotic replicative CMG helicase"

**For**

**DNA**

The DNAs used for these studies were the following oligonucleotides (Integrated DNA Technologies) of: a 45-mer lagging-strand oligo, (5'-GGCAGGCAGGCAGGCACACACTCTCCAATTA/iBiodT/CACTTCCTACTCTA-3') and a 70-mer leading-strand oligo (5'-TAGAGTAGGAAGTGA/iBiodT/AATTGGAGAGTGTGTTTTTTTTTTTTTTTTTTTTTTTTTTTTTTTTTTT*T*T*T*T*T-3'). The asterisks are residues containing a phosphothio linkage. The two oligos were annealed in equimolar amounts by heating to 90°C followed by slow (1 hour) cooling to room temperature. The hybrid was purified from a 8% native PAGE then a 1 molar equivalent of streptavidin was added, enabling a single streptavidin to cross-link the two diametrically opposed biotins.

**Proteins**

Mcm10 and CMG were purified as previously described (Langston et al., 2017), with the following modifications. The Mcm10 contained a N-terminal hexahistidine tag and a C-terminal 3X FLAG tag. Briefly, 48 L E. coli cells carrying the Mcm10 T7 based E. coli expression vector were grown to OD 0.6 at 37 $^{o}$C, then cooled to 15 $^{o}$C and induced upon adding IPTG for an additional 8 h at 15 $^{o}$C. The cells were harvested by slow speed centrifugation and lysed using a continuous flow high speed homogenizer. Cell debris was removed by centrifugation and applied to a 10 ml Chelating Sepharose Fast Flow column (GE Healthcare) charged with 50 mM $NiSO_4$ in Buffer A (20 mM Tris-Cl pH 7.9, 5 mM imidazole, 500 mM NaCl, 0.01% NP-40). The column was washed with Buffer A, then eluted with 375 mM imidazole in Buffer A. The eluated material was applied to a 6 ml anti-FLAG M2 affinity gel (Sigma) equilibrated in Buffer B (20 mM Tris-Cl pH 7.5, 10% glycerol, 500 mM NaCl, 1 mM DTT, 1 mM $MgCl_2$, 0.01% NP-40) and then washed with 20 column volumes of Buffer B before eluting with Buffer B containing 0.2 mg/ml FLAG peptide (EZ Biolab, Carmel, Indiana USA) using two 6 ml pulses of 20 min each and collecting 1.5 ml fractions. Eluted fractions were analyzed by SDS-PAGE, protein concentration was determined using Bradford Protein Stain (Sigma) wtih BSA as a standard. Proteins were then aliquoted, snap frozen in liquid nitrogen and stored at -80 $^{o}$C.

The CMG-Mcm10 complex was reconstituted by mixing 765 pmol CMG with 3.1 nmol Mcm10 in 0.7 ml Buffer C (10 mM Tris-Cl pH 7.5, 200 mM KCl, 2mM DTT, 2 mM $MgCl_2$) for 30 minutes on ice. The mixture was then applied to a 0.1 ml MonoQ column, equilibrated in Buffer C. The column was washed using the same buffer and eluted with a 2.5 ml gradient of Buffer C from 0.2 M KCl to 0.6 M KCl. Fractions of 0.1 ml were collected, analysed by Bradford and SDS-PAGE, pooled and dialyzed against 50 mM K-glutamate in 25 mM Tris-acetate pH 7.5, 2 mM Mg acetate, and 1 mM DTT. Dialysed material was analyzed again by Bradford stain for protein concentation , then aliquoted, snap frozen in liquid nitrogen and stored at -80 $^{o}$C.

**Sample preparation for CryoEM**

The replication fork bound to streptavidin (10 µM final) was added to a solution containing 1.2 mg/ml CMGM in 20 mM Tris acetate pH 7.5, 40 mM K-glutamate, 40 mM KCl, 1 mM DTT, 2 mM Mg acetate, along with 0.1 mM ATP. The sample was incubated for 5 min at room temperature, separated into 10 ml aliquots, snap frozen in liquid nitrogen, and stored at -80 $^{o}$C. Samples were applied to cryo-EM grids immediately upon thawing.

**Cryo-EM**

To prepare EM grids, 3 µl of CMG-Mcm10-bio-DNA-SA sample was applied, at a final concentration of ~ 1.0 mg/ml, to C-flat 1.2/1/3 holey carbon grids, treated by glow-discharge before use. Grids were then incubated for 10 s at 6 °C and 90% humidity and blotted for 3 s and plunged into liquid ethane using a Thermo Fisher (TF) Vitrobot IV. Grids were loaded into a TF Titian Krios electron microscope and images at 300 keV were collected automatically using low-dose mode at a magnification of ×130,000 and a pixel size of 1.029 Å per pixel. A Gatan K2 summit direct electron detector was used for image recording with a defocus range from 1.5 to 3.5 µm under super-resolution mode. The dose rate was 10 electrons per Å$^2$ per second and total exposure time was 8 s. The total does was divided into a 40 frame movies and each frame was exposed for 0.2 s.

**Image processing and 3D reconstruction**

Over 6,000 raw movie micrographs were collected. Firstly, all the movie frames were aligned and superimposed by Motioncorr2. Contrast transfer function parameters of each aligned micrograph were calculated with CTFFIND4. We manually picked about 5, 000 particles from different views to generate several 2D averages used as a template for automatic particle picking. Automatic particle picking was then performed for the whole data set. About 718,903 particles were initially picked. These were then sorted according to the similarity to the 2D reference; the bottom 10% particles that had very low z-scores were deleted from the particle pool. The 2D classification of all the remaining particles was performed and particles in bad classes (i.e. no CMG structure observed) were removed. 387,023 good particles (CMG structure was observed) were kept for the following 3D classification. We derived five 3D models from the dataset: two models were identified having fork DNA and these associated particles were combined for further refinement; the other three models showed no DNA inside or were distorted and those particles were discarded. A total of 162,550 particles were used for the final refinement, leading to the final 3D map with an estimated average resolution of 3.9 Å. The resolution estimations were based on gold-standard Fourier shell correlation calculations to avoid over-fitting and reported resolutions were based on the FSC = 0.143 criterion. All the density maps were corrected for the modulation transfer function of the detector and sharpened by applying negative B-factors. All the steps mentioned above, including particles autopicking, 2D classification, 3D classification, 3D refinement, postprocess, were done in Relion-2.0. Local resolution was estimated using ResMap. These steps are illustrated in Figs. S1-S3.

### Supplemental Figures

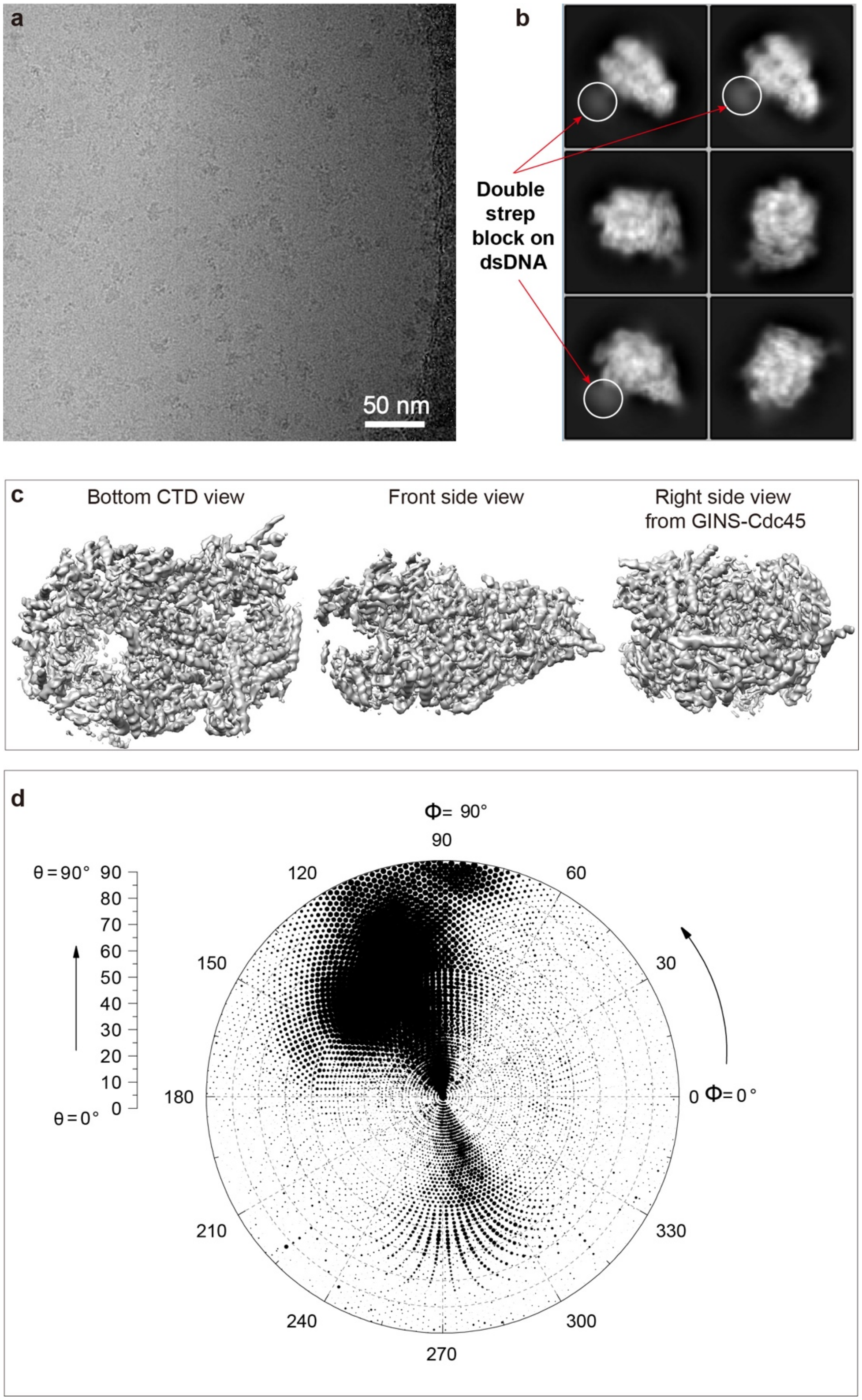


**Figure S1. Cryo-EM of CMG bound to double-streptavidin blocked DNA fork in the presence of Mcm10 and ATP.** (**a**) A representative raw micrograph. (**b**) 2D class averages. (**c**) 3D map in different views. (**d**) Euler angle distribution of the particles in the final reconstruction.

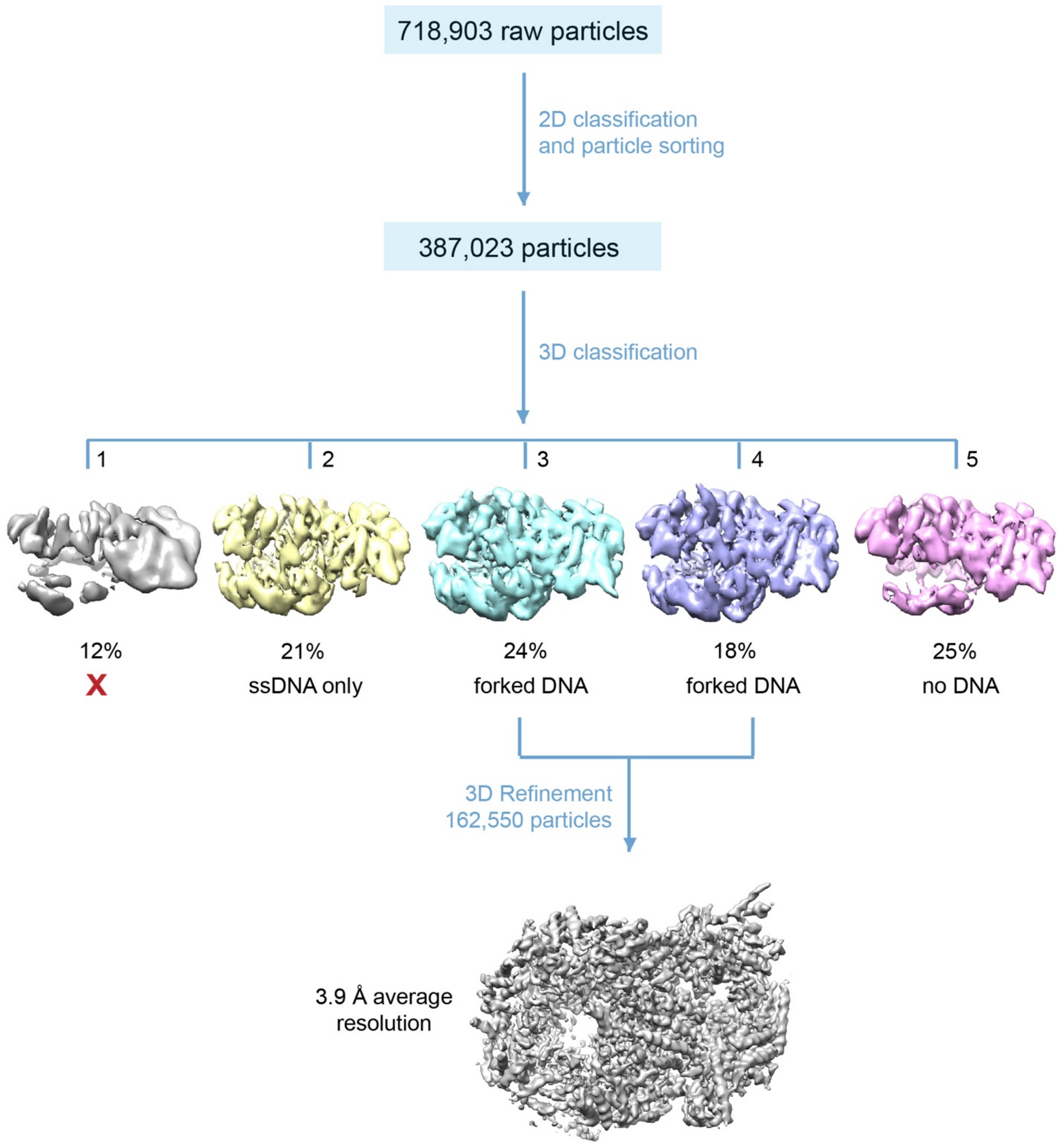


**Figure S2. Image processing and 3D reconstruction procedure.** Raw particles from auto-picking were classified and those with clear class averages were selected for 3D classification. Among five 3D classes, only two contained forked DNA density. These two classes were selected for final refinement and 3D reconstruction, leading to the 3.9-Å resolution 3D map.

a

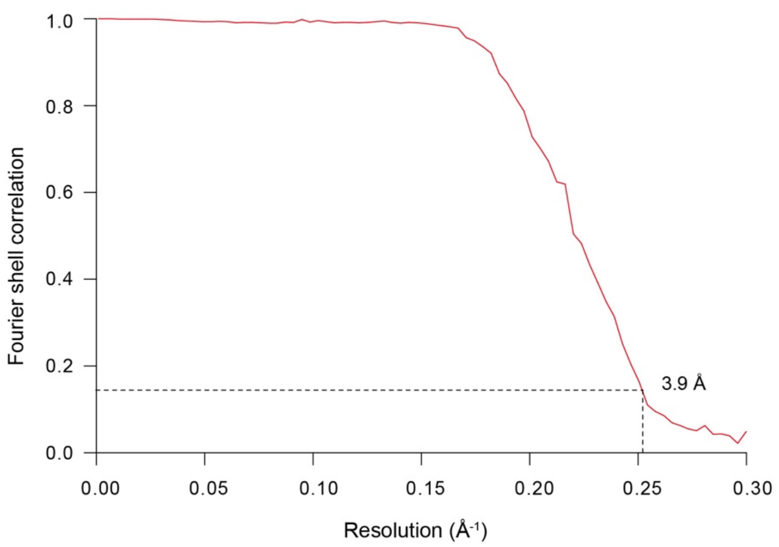


b

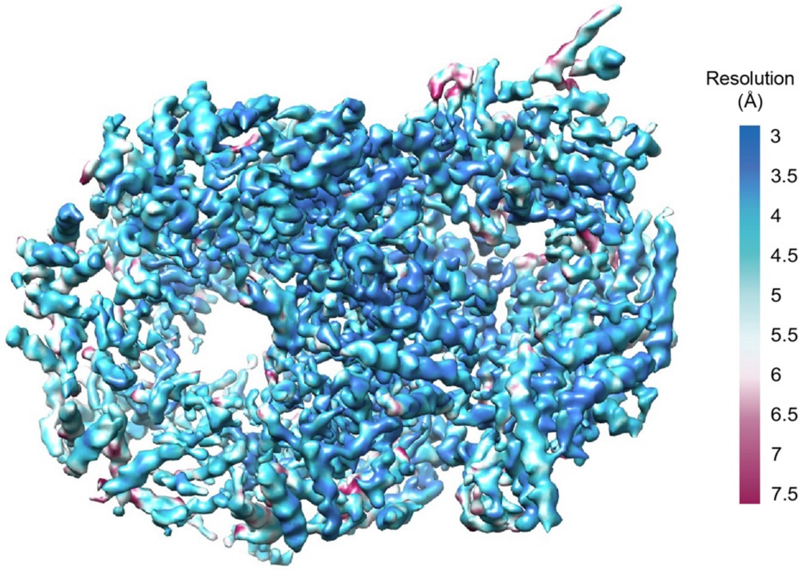


**Figure S3. Resolution estimation of the final 3D map of the CMG-forked DNA.** (**a**) Gold standard Fourier shell correlation indicated an average resolution of 3.9 Å. (**b**) Local resolution map. Note that the parental dsDNA density is invisible at the raised display threshold.

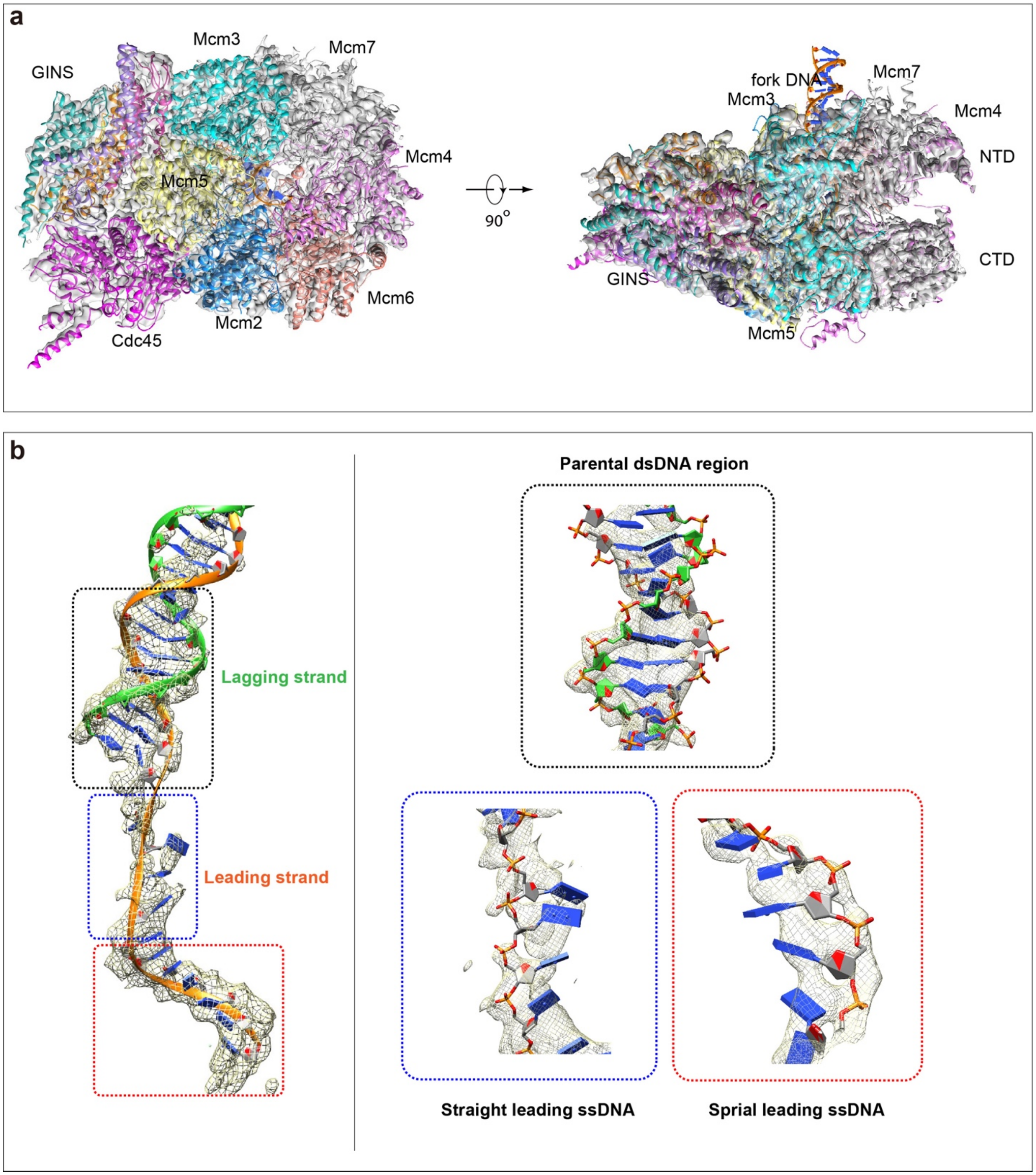


**Figure S4. Surface-rendered cryo-EM 3D map of the CMG-forked DNA superimposed with the atomic model.** (**a**) Overall view of the cryo-EM map in two orthogonal views. Note that the parental dsDNA density outside CMM is invisible at this raised display threshold. (**b**) Cryo-EM density of the forked DNA. Zoomed views of the parental DNA region, the linear leading ssDNA region, and the spiral leading ssDNA region, are shown to the right.

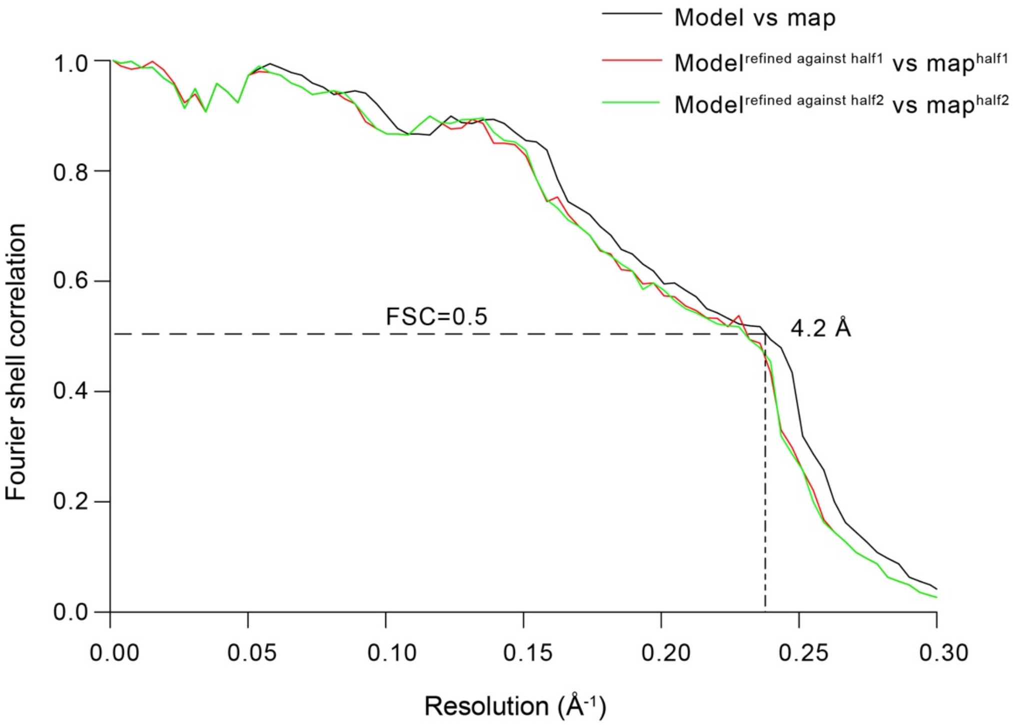


**Figure S5. Map and atomic model correlation validation.** The atomic model is estimated to have an average resolution of 4.2 Å.
